# Morphoregulatory ADD3 underlies glioblastoma growth and formation of tumor-tumor connections

**DOI:** 10.1101/2024.03.26.586725

**Authors:** Carlotta Barelli, Flaminia Kaluthantrige Don, Raffaele M. Iannuzzi, Ilaria Bertani, Isabella Osei, Simona Sorrentino, Giulia Villa, Viktoria Sokolova, Francesco Iorio, Nereo Kalebic

## Abstract

Glioblastoma is a major unmet clinical need characterized by striking inter- and intra-tumoral heterogeneity and a population of glioblastoma stem cells (GSCs), conferring aggressiveness and therapy resistance. GSCs communicate through a network of tumor-tumor connections (TTCs), including nanotubes and microtubes, promoting tumor progression. However, very little is known about the mechanisms underlying TTC formation and overall GSC morphology. As GSCs closely resemble neural progenitor cells during neurodevelopment, we hypothesised that GSCs’ morphological features affect tumour progression. We identified GSC morphology as a new layer of tumoral heterogeneity with important consequences on GSC proliferation. Strikingly, we showed that the neurodevelopmental morphoregulator ADD3, is sufficient and necessary for maintaining proper GSC morphology, TTC abundance and cell cycle progression as well as required for cell survival. Remarkably, both the effects on cell morphology and proliferation depend on the stability of actin cytoskeleton. Hence, cell morphology and its regulators play a key role in tumor progression by mediating cell-cell communication. We thus propose that GSC morphological heterogeneity holds the potential to identify new therapeutic targets and diagnostic markers.

## Introduction

Glioblastoma (GBM) is the most aggressive and common form of primary brain malignancy in adults and an unmet clinical need (Tran and Rosenthal, 2010). Its high chance of relapse is largely due to its striking inter- and intra-tumoral heterogeneity along with its infiltration into the healthy brain parenchyma (Garofano et al., 2021; Petrecca et al., 2013; Spiteri et al., 2019). Cellular interactions between GBM cells and the microenvironment were shown to be important to maintain the aggressive character of the tumor (Osswald et al., 2016; Pinto et al., 2020; Yabo et al., 2022). GBM cells form intercellular networks via two main types of tumor cell-tumor cell connections (TTCs): tunneling nanotubes (TNTs) and tumor microtubes (TMs) (Pinto et al., 2020; Venkataramani et al., 2022a; Zurzolo, 2021). Through these connections, cancer cells form a multicellular network which affects GBM proliferation (Osswald et al., 2015; Ratliff et al., 2023), invasion (Lu et al., 2017; Lu et al., 2019; Osswald et al., 2015; Venkataramani et al., 2022b) and therapy resistance (Hekmatshoar et al., 2018; Kolba et al., 2019; Osswald et al., 2015; Weil et al., 2017). Despite such important role of cellular protrusions in GBM, little is known about the morphological heterogeneity of GBM cells, the molecules underlying it and its role in cell proliferation.

Given the striking similarities between neurodevelopment and GBM progression, neural progenitor cells could offer key insights into the molecular and cellular underpinnings of GBM cell morphology and its role in cancer progression. Moreover, a specific type of GBM cells, known as glioblastoma stem cells (GSCs), which confer aggressiveness and therapy resistance to the tumor (Azzarelli et al., 2018; Neftel et al., 2019), shows remarkable similarities with a population of neural progenitor cells called basal or outer radial glia (bRG or oRG), a key cell type underlying fetal development of the human cortex (Fietz et al., 2010; Hansen et al., 2010; Reillo et al., 2011). Not only do GSCs show transcriptomic signatures of bRG (Bhaduri et al., 2020; Couturier et al., 2020), but they also undergo a characteristic type of cell movement, called mitotic somal translocation (MST), previously reported only in fetal bRG (Bhaduri et al., 2020; Hansen et al., 2010; LaMonica et al., 2013). In bRG, cell morphology was shown to have an important role in underlying cell proliferation, migration and MST (Del-Valle-Anton and Borrell, 2022; Kalebic and Huttner, 2020; Molnar et al., 2019; Ostrem et al., 2017; Taverna et al., 2014). In fact, different bRG morphotypes were identified (Betizeau et al., 2013; Kalebic et al., 2019; Reillo et al., 2017) and increased morphological complexity has been linked to a greater proliferative potential (Kalebic et al., 2019). Considering such role of cell morphology in neurodevelopment and the presence of tumor microtubes in GBM, we hypothesized that morphological complexity affects GBM progression.

Here we identified adducin-γ (ADD3), an actin-associated protein known to control bRG morphology and proliferation, as a putative master morphoregulator of GSCs. We next investigated the morphological heterogeneity of GSCs in different GBM cell lines and found that they exist in four morphotypes, similar to neural progenitors in the developing brain. We demonstrated that ADD3 regulates the morphology of GSCs by inducing their elongation, branching and the formation of tumor microtubes. We further showed that the effect of ADD3 on cell morphology are necessary for cell survival and proliferation. Hence, we described cell morphology as a new layer of heterogeneity in GBM and identified morphoregulatory proteins as potential targets to tackle GBM progression.

## Results

### Identification of morphoregulatory adducin-γ (ADD3) in GBM

Considering the resemblance between GSCs and bRG, we first sought to identify genes that might govern the GSC morphology by mining datasets of morphoregulators in fetal bRG. We combined previously published transcriptional (Fietz et al., 2012) and proteomic (Kalebic et al., 2019) analyses and identified 45 morpho-regulatory genes whose expression is enriched in bRG *vs*. other cell types of the developing brain. We next intersected this list with a published list of genes expressed in glioblastoma (Bhaduri et al., 2020) (Figure 1A). Among the 30 identified genes, the adducin family was prominently present (Fisher exact test *p-value* = 3.34 x 10^-9^) with all its three members (Figure 1B, B’).

**Figure 1.**
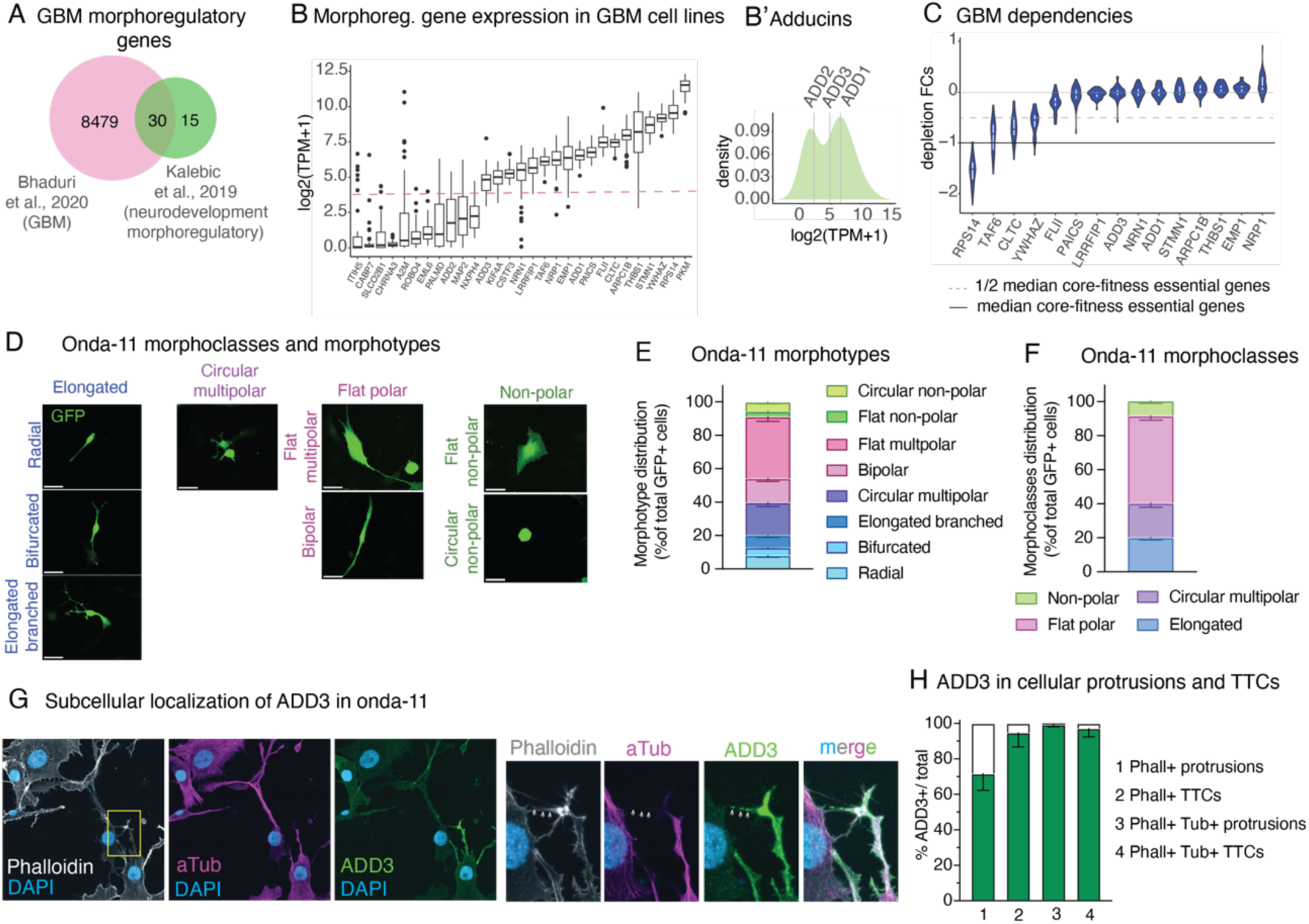
Onda 11 GBM cells show morphological heterogeneity and are dependent on ADD3, a neurodevelopmental morphoregulator localised in GBM cell protrusions and TTCs. (A-C) Computational identification of ADD3 as a neurodevelopmental morphoregulator with a putative role in GBM progression. Data come from (Bhaduri et al., 2020) and (Kalebic et al., 2019) (A) and Broad DepMap 22Q2 version and Sanger cell model passport (B and C) (A) Intersection between a list of 8509 differentially expressed genes in primary glioblastoma tumors and 45 neurodevelopmental morphoregulatory genes, resulting in 30 shared genes. (B) Log2(TPM+1) expression levels of the resulted gene list (29/30) averaged across 48 annotated glioblastoma cell lines from showing bimodal distribution (dashed line). (B’) Density plot of the average expression levels of the adducin family of genes indicating the estimated density with superimposed average expression levels of adducins. (C) Dependency of GBM cell lines (depletion fold change (FC) distribution upon CRISPR/Cas9 targeting) on the 15 highly expressed non-core-fitness genes from panel (B). (D) Onda 11 GSCs were transfected with GFP and their cell morphology was analyzed 72 h later. MIP of 12 planes. Four different morphoclasses listed at the top of the images (elongated, circular multipolar, flat polar, non-polar) are further divided into eight morphotypes annotated on the left of the images (radial, bifurcated, elongated branched, circular multipolar, flat multipolar, bipolar, flat non-polar, circular non-polar). Scale bars; 10µm. (E-F) Analysis of Onda-11 morphology using GFP signal, 72 h after transfection, showing their morphological heterogeneity. Distribution of the 8 morphotypes (E) grouped into 4 morphoclasses (F). Mean of 8 independent transfections. Error bars, SEM. (G-H) ADD3 is expressed in cellular protrusions and tumor cell-tumor cell connections (TTCs) of Onda11 GSCs. (G) IF staining for actin (Phalloidin, white), microtubules (alpha-Tubulin, magenta), ADD3 (green) along with DAPI staining (blue), MIP of 12 planes. Image size 188.61 x 188.61 µm (top); 35.69 x 53.14 µm (bottom). (H) Quantification of the expression of ADD3 in Onda11 GSC protrusions and microtubes. Error bar, SD; n = 3 independent cell cultures.

Adducins are morphoregulatory proteins involved in the assembly of the actin-spectrin network and are implicated in growth of cell protrusions, membrane trafficking and providing mechanical stability to plasma membrane (Baines, 2010; Kiang and Leung, 2018; Kiang et al., 2020; Lou et al., 2013). Taking advantage of data from the Cell Model Passports (van der Meer et al., 2019) and the Cancer Dependency Maps (Behan et al., 2019; Pacini et al., 2021; Tsherniak et al., 2017), we excluded genes that were not expressed at the basal level in a panel of commercially available and multi-omically characterised GBM cell lines (Figure 1B) and that are core-fitness essential genes (Vinceti et al., 2021) (Figure S1A) shortlisting a set of 15 candidate genes (Figure 1C). Of the 3 adducins, 2 (ADD1 and ADD3) were in this list, with adducin-γ (ADD3) showing a strong and seemingly context-specific, essentiality (Figure 1C). The role of ADD3 in glioblastoma is not clear, as it has been reported to both promote and reduce tumor growth and invasiveness (Kiang et al., 2020; Rani et al., 2013). Furthermore, ADD3 has been associated with temozolomide resistance (Poon et al., 2015), glioma progression (Rani et al., 2013; van den Boom et al., 2003) and reduced glioma cell motility (Mariani et al., 2001). Strikingly, we have previously shown that ADD3 is required for the correct morphology of human basal progenitors and that its depletion results in a reduction of their proliferation (Kalebic et al., 2019).

### Morphological heterogeneity of GSCs and subcellular localization of ADD3

Analysis of Cancer Dependency Map datasets revealed that the glioblastoma cell line Onda-11 exhibits the strongest dependency on ADD3 (scaled depletion fold-change upon CRISPR-Cas9 targeting = −0.59, with −1 indicating the median depletion fold-change of strongly essential core-fitness genes, such as ribosomal protein genes Figure 1C and S1B). To promote stemness of Onda-11 cells, we maintained them in serum-free culture conditions and confirmed their stem-like features by immunofluorescence for nestin, SOX2, L1CAM, OCT4, GFAP and CD44 (Figure S1C-I).

Upon transfection with GFP we examined the morphology of Onda-11 GSCs, and found a remarkable heterogeneity identifying eight morphotypes, which we grouped into four principal morphoclasses: non-polar, flat polar, circular multipolar and elongated (Figure 1D-F). Morphologically these cells were reminiscent of neural progenitor cells during cortical development (Kalebic and Huttner, 2020). Specifically, elongated morphoclass (radial and bifurcated morphotypes) and bipolar cells of the flat polar morphoclass, morphologically resemble morphotypes of bRG, whereas circular multipolar cells resemble multipolar basal progenitors (Kalebic et al., 2019). Instead, flat multipolar and flat non-polar GSCs do not seem to have a corresponding developmental morphotypes and likely arise during tumorigenesis. This suggest that, in addition to the molecular and cell behavioral features (Bhaduri et al., 2020), GSCs also recapitulate the morphological features of embryonic neural progenitors, in particular of bRG. We further confirmed the existence of the four morphoclasses in U87-MG glioblastoma cell line (see Figure S3D).

We examined the subcellular localization of ADD3 in Onda-11 GSCs by confocal microscopy. We observed that ADD3 readily localizes to the proximity of plasma membrane, to cellular protrusions and, specifically, TTCs (Figure 1G). Whereas ADD3 was enriched in protrusions that contained both microtubules and actin, in TTCs it was present irrespectively of whether they contained actin only or actin and microtubules (Figure 1H). Considering the morphological heterogeneity of GSCs and the localization of ADD3 to cellular protrusions, we next sought to examine the potential ability of ADD3 to affect GSC morphology and its role in glioblastoma growth.

### ADD3 is sufficient and required to control the number of protrusions and elongation of GSCs

We transfected Onda-11 GSCs with ADD3-over-expressing (ADD3 OE) and control plasmids along with GFP, to visualize cell shape, and performed a morphological analysis three days following transfection (Figure 2A and S2A). ADD3 OE led to an altered distribution of morphoclasses with a marked increase in the proportion of elongated cells at the expense of the other three morphoclasses (Figure 2B). To examine various features of the cell morphology in a quantitative manner, we established a machine learning-assisted pipeline for the automatic segmentation and analysis of microscopy images (Figure 2C). Employing this pipeline to examine the effects of ADD3 OE, we observed a striking increase in the number of cellular protrusions (Figure 2D), both primary protrusions that grow directly from the cell body (Figure 2E), and all protrusions, which include also secondary and other higher-order protrusions, compared to the control. This was accompanied by an increase in both the average and the maximum length of cell protrusions (Figure 2F, G), which was confirmed also by the Scholl analysis (Figure S2B), and by an increase in protrusion branching (Figure 2H). Together, this suggestes that ADD3 promotes both the formation and the growth of new protrusions. Besides, such an increase in cellular protrusions also enlarged cell perimeter and area (Figure 2I, J). Finally, the overall shape of ADD3-over expressing cells became more elongated, as their major axis was significantly longer than in the control cells, whereas the length of the minor axis was not affected (Figure 2K, L). Accordingly, cell eccentricity was increased, indicating a more elliptic and elongated shape as opposed to circular (Figure 2M).

**Figure 2.**
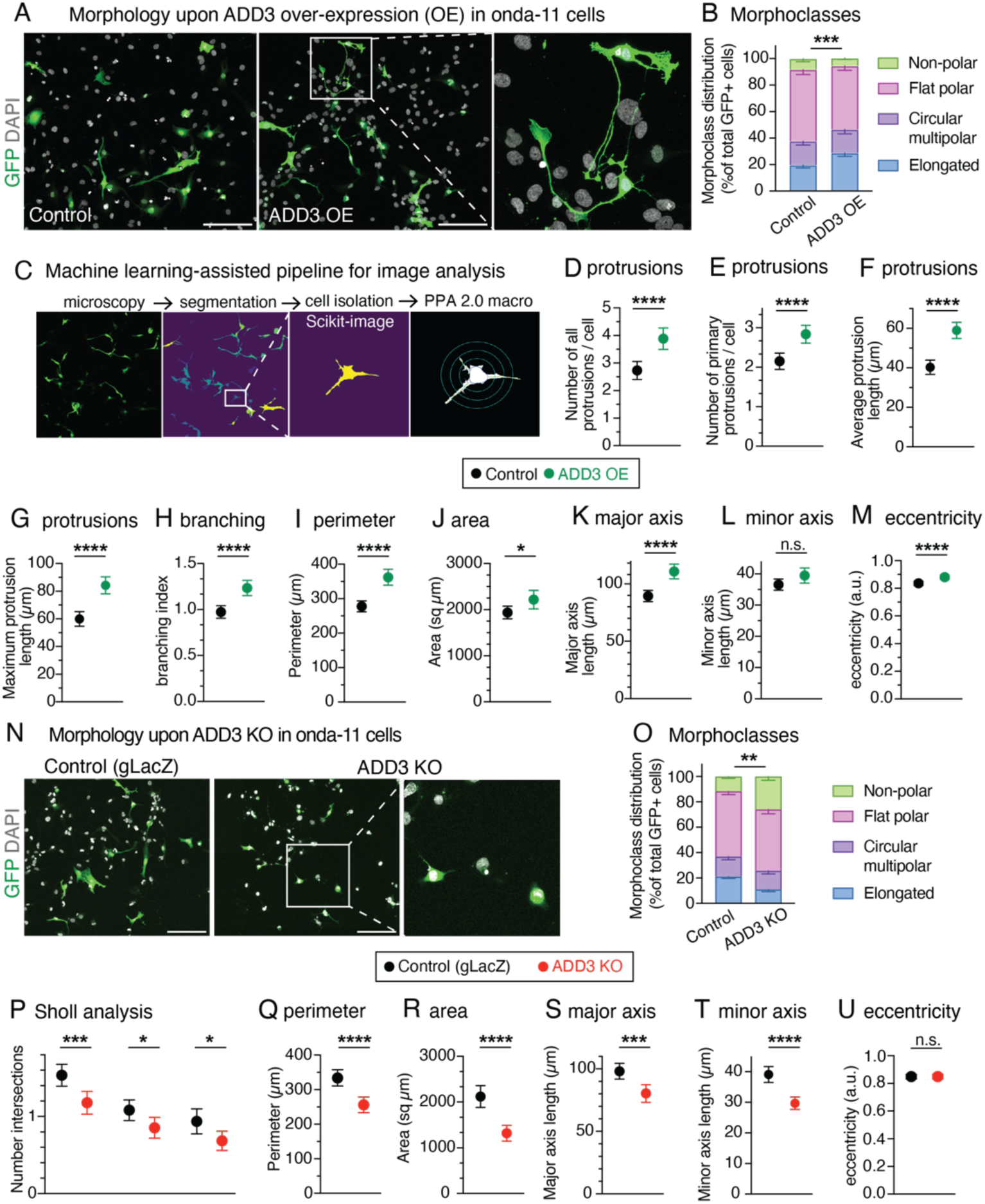
ADD3 regulates Onda 11 GSC morphology and protrusion number. (A-M) ADD3 overexpression promotes cell elongation and protrusion abundance. Onda 11 cells were transfected either with GFP and ADD3 overexpressing plasmids (ADD3 OE) or with GFP and empty vector (control) and their morphology was analyzed. (A) Representative examples of GFP+ (green) Onda 11 cell morphology in control (left) and ADD3 OE (center). Scale bar, 200 µm. A close-up of elongated cells upon ADD3 OE (right, image width 250 µm). MIP of 12 planes. (B) Distribution of the 4 morphoclasses in control and ADD3 OE Onda 11 GSCs. (C) Schematics of the pipeline for automated cell segmentation and morphological analysis of cells. GFP+ cells from confocal microscopy images (MIPs of 25 planes) are segmented in CellPose and single cells are isolated to carry out morphological analysis in Python and Fiji using PPA 2.0 macro. (D-M) Number of total (D) and primary (E) cell protrusions, average (F) and maximum (G) protrusion length, branching index (H), perimeter (I), area (J), major (K) and minor (L) axis length and eccentricity (M) upon ADD3 OE vs. Control, calculated as described in (C). (N-U) ADD3 KO reduces protrusion abundance and induces cell shrinkage. Onda 11 cells were transfected either with ADD3 KO plasmid or with gLacZ KO plasmid as control and their morphology was analyzed. (N) Representative examples of GFP+ (green) Onda 11 cell morphology in control (left) and ADD3 KO (center). Scale bar, 200 µm. A close-up of cells upon ADD3 KO (right, image width 300 µm). MIP of 12 planes. (O) Distribution of the 4 morphoclasses in control and ADD3 KO Onda 11 GSCs. (P-U) Sholl analysis (P), perimeter (Q), area (R), major (S) and minor (T) axis length and eccentricity (U) upon ADD3 KO vs. Control, calculated as described in (C). (B, D-M, O-U) Mean of 4 (B, D-M) and 8 (O, P-U) independent transfections. Total number of cells scored: 328 (ADD3 OE) and 397 (control) (D-M); 317 (KO and control) (P-U). Error bars, SEM (B, O), 95% CI (D-M, P-U); *, P<0.05; **, P<0.01; ***, P<0.001; ****, P<0.0001; n.s. not statistically significant; two-way ANOVA with Sidak’s post hoc tests (B, O), Student’s t-test (D-M, P-U).

We next examined if ADD3 was required to maintain the correct Onda-11 morphology. We performed a CRISPR/Cas9-mediated knock-out (KO) of ADD3 and confirmed its efficiency by both immunoblot and immunofluorescence three days after transfection (Figure S2C, D). Inspection of the Onda-11 morphology upon ADD3 KO showed altered distribution of morphoclasses with an apparent reduction in the proportion of the elongated cells and a relative increase in the non-polar cells (Figure 2N, O). Consistent with this and opposite to the effects of the over expression, ADD3 KO resulted in the reduction of the number of protrusions, their length, branching index, cell perimeter and area (Figure 2P-R, S2E-J). This was accompanied by a reduction in both major and minor axis length (Figure 2S, T), but did not result in a statistically significant reduction in cell eccentricity (Figure 2U).

We examined if the above effects of ADD3 on cell morphology are pertinent to other glioblastoma cell lines. We performed ADD3 KO in U87-MG glioblastoma line and H4 neuroglioma line (Figure S3A, B, E, F). Whereas U87-MG showed strong morphological heterogeneity, which was comparable to Onda-11, H4 cells exhibited rather uniform morphologies (Figure S3C, D, G). Accordingly, KO of ADD3 resulted in a change in morphotype distribution in U87-MG, but not H4 cells (Figure S3D, G). Similarly to Onda-11, ADD3 KO in U87-MG cells resulted in an increase in non-polar cells at the expense of elongated ones. This suggests that the effects of ADD3 are pertinent to other glioblastoma cell lines, particularly those that exhibit a heterogeneous cell morphology.

Taken together, these analyses show that ADD3 is both sufficient and required to maintain correct cell morphology, including the correct number and length of cellular protrusions, their branching, cell size and elongation. Instead, ADD3 is sufficient to increase cell eccentricity, while its KO resulted in cell shrinkage without modifying the eccentricity.

### ADD3 promotes morphological transitions during interphase

Considering the above change in the distribution of morphoclasses we sought to examine potential transitions between GSC morphoclasses using time-lapse microscopy. Two days following transfection with GFP and ADD3 or control plasmids, Onda-11 GSCs were imaged for 60 hours. We first focused on the morphological dynamics during interphase and observed that Onda-11 GSCs only rarely undergo a transition between morphoclasses (Figure 3A, B and Movie S1). In fact, only the nonpolar cells exhibited morphological dynamics in interphase (Figure 3C, S4A). ADD3 was however able to promote such morphoclass transitions (Figure 3A, B) with an increase in transitions into elongated cells (Figure 3A, C and Movie S2 and compare to Movie S1).

**Figure 3.**
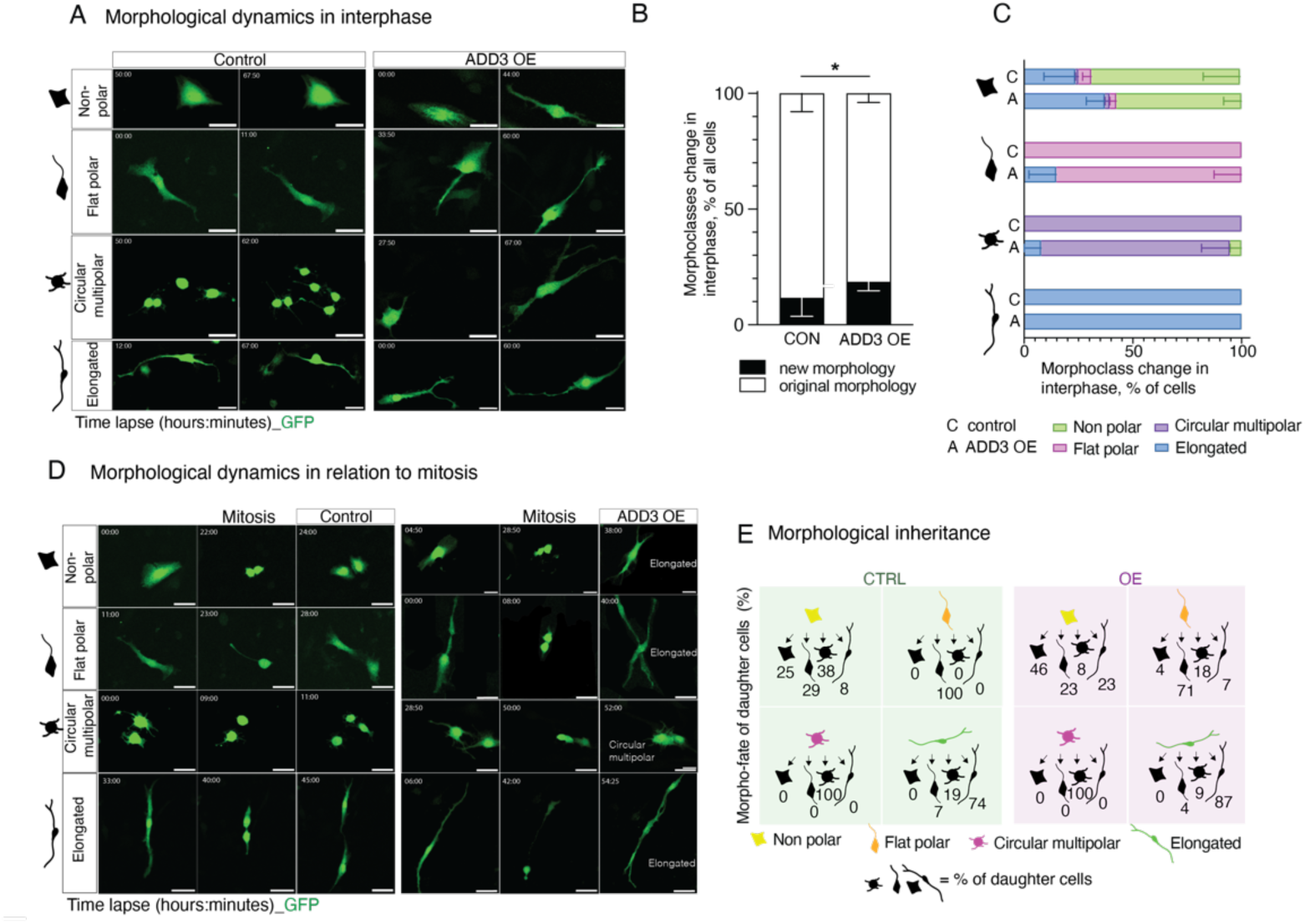
ADD3 promotes morphological transitions in interphase. Onda 11 cells were transfected either with GFP and ADD3 overexpressing plasmids (ADD3 OE) or with GFP and empty vector (control) and their morphological dynamics were analyzed by live imaging in interphase (A-C) and in relation to mitosis (D, E). (A) Examples of the morphological dynamics in interphase of the 4 morphoclasses upon ADD3 OE vs Control. Note the increased elongation of the ADD3 OE cells. The time lapse indicated in the upper left corner of the images. Scale bars, 50 µm. (B) Quantification of morphological changes in interphase. Note the increase in acquisition of new morphology upon ADD3 OE. Mean of 4 independent transfections. Error bars, SD; *, P<0.05; two-way ANOVA with Bonferroni post hoc tests. (C) Quantification of morphological transitions in interphase for each morphoclass. Mean of 3 independent transfections. Error bars, SEM. (D) Examples of the morphological dynamics in relation to mitosis of the 4 morphoclasses showing the time point pre-(left), during (middle) and post-mitosis (right) upon ADD3 OE vs Control. The time lapse indicated in the upper left corner of the images. Scale bars, 50 µm. (E) Schematic representation of the morphological inheritance shown as the percentage of morphoclass progeny for each mother morphotype. Number of mother cells (control, ADD3 OE): nonpolar (12, 13), flat polar (16, 16), circular multipolar (30, 5), elongated (15, 29). Data come from 4 independent transfections. See also Supplemental Movies S1-5.

As mitosis involves characteristic morphological changes, we specifically examined the inheritance of the mother cell morphology upon the cell division. During the duration of the live imaging, around 40% of cells underwent mitosis (Figure S4B). In control the mother cell morphology was generally inherited by both daughter cells (Figure 3D, E, S4C). Among the four morphoclasses, non-polar cells again displayed the greatest number of transitions (Figure 3E and Movie S4). Differently to what observed in interphase (Figure 3B), ADD3 OE was not able to alter the frequency of morphoclass transitions in mitosis (Figure S4C and Movie S3). However, upon ADD3 OE we observed (1) morphological transitions of the progeny of flat polar dividing cells (Figure 3D, E and S4D) as well as (2) a subtle increase in the elongated progeny of the diving cells (Figure 3E, S4D and Movie S5). Both effects were similar to what described above for the interphase. Interestingly, in a subset of elongated cells, we observed an MST-like behavior, which however did not seem to be regulated by the over expression of ADD3 (Figure S4E).

Taken together, these data show that the morphoclass identity is largely conserved both in interphase and in relation to mitosis. The morphological heterogeneity instead seems to be principally generated by the morphological dynamics of non-polar cells in both interphase and mitosis. ADD3 over expression led to an increase in transitions from all morphoclasses into elongated cells, which is consistent with the increase in the proportion of elongated cells described above (Figure 2). Finally, while this effect was mild in mitosis, it led to a marked increase in elongated cells during interphase.

### ADD3 controls Onda-11 GSCs proliferation and survival

In light of the (1) effects of ADD3 on GSC morphology (Figure 2) and the previous data showing that (2) ADD3 underlies progenitor morphology and proliferation during cortical development (Kalebic et al., 2019), we sought to examine the putative effects of ADD3 on the proliferation of Onda-11 GSCs. We first examined the expression pattern of Ki67, a marker of cell proliferation, and categorized cells in three phases of the cell cycle (Figure S5A). Upon ADD3

OE we detected a relative increase in the proportion of cells in G0 and early G1 phases (Figure 4A, B). This led to a marked reduction in the proportion of cells in late G2 and M-phase, which was confirmed also by immunostaining for a mitotic marker, phospho-vimentin (pVim, Figure S5B-C). Investigating the Ki67 expression pattern across the four morphoclasses, revealed the strongest effect in circular multipolar cells and less prominently in elongated and nonpolar cells (Figure 4C). Upon EdU treatment of Onda-11 GSCs, we detected no difference in the proportion of cells in S-phase (Figure S5D-E), suggesting that the principal effects of ADD3 OE on GSC proliferation are related to G0/early G1 phases and mitosis.

**Figure 4.**
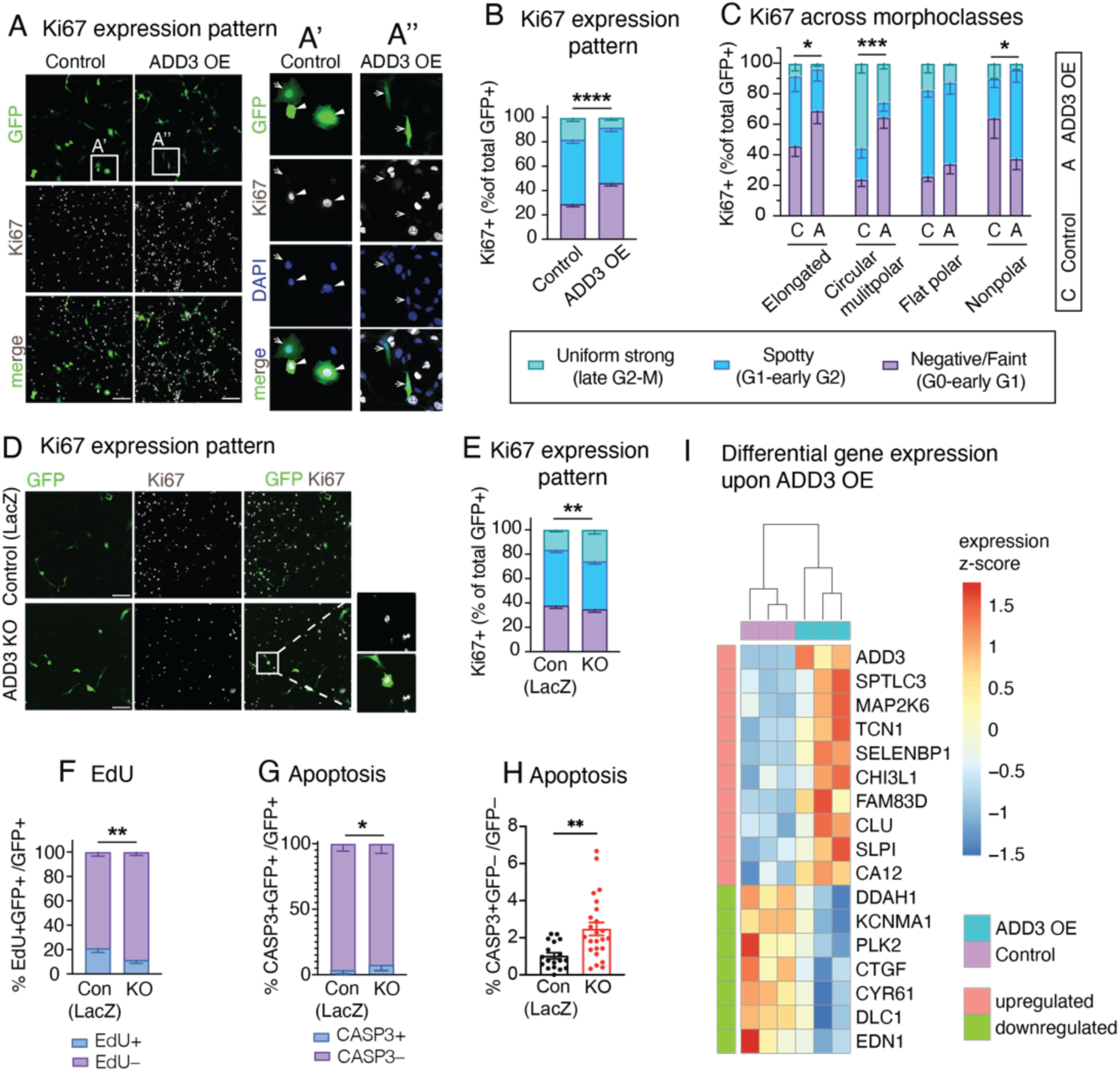
ADD3 regulates Onda 11 GSC proliferation and survival. (A-C, E, F) Effects of ADD3 OE (A-D) and KO (E, F) on cell proliferation 72 h after transfection, analyzed by IF for the expression pattern of Ki67, which is indicative of different phases of the cell cycle: uniform strong (late G2-M, green), spotty (G1-early G2, blue), negative/faint (G0-early G1, violet). MIPs of 12 planes are used to analyse Ki67 pattern of expression. (A) Representative images of IF for Ki67 (white) along with DAPI staining (blue) in GFP+ cells (green). Close-ups of GFP+ control (A’) and ADD3 OE cells (A’’). Arrows, Ki67 negative/faint; arrowheads, Ki67 spotty/uniform strong. Note the negative/faint Ki67 expression upon ADD3 OE. (B, C) Distribution of the three Ki67 expression patterns in control and ADD3 OE Onda 11 GSCs in the whole population (B) and across morphoclasses (C). (D) Representative images of IF for Ki67 (white) along with DAPI staining (blue) in GFP+ cells (green). Close-ups of GFP+ ADD3 KO cells. (E) Distribution of the three Ki67 expression patterns in control and ADD3 KO Onda 11 GSCs. Note that ADD3 KO increases the percentage of Onda 11 GSCs in late G2/M. (F) Effects of ADD3 KO on cell proliferation 72 h after transfection, analyzed by EdU treatment (4 h) and microscopy. Distribution of EdU+ and EdU-GFP+ Onda 11 GSCs upon ADD3 KO. (G, H) Effects of ADD3 KO on cell apoptosis 72 h after transfection, analyzed by IF for cleaved caspase-3 (CASP3) in GFP+ transfected cells (H) and GFP– cells (I). Note the increase in cell apoptosis upon ADD3 KO in both transfected and surrounding cells. (I) Differentially expressed genes from contrasting bulk RNA-seq profiles of ADD3 OE Onda-11 vs Control, 72 h after transfection. Z-scores of differentially expressed genes (absolute log FC ≥ 0.5 and adjusted p.value < 0.05) are grouped row-wise according to differential expression sign, with samples hierarchically clustered based on Euclidean similarity. (A, E) Scale bars, 200 µm (A, E). Image width, 232 µm (A’, A’’); 200 µm (E, insets). (B, C, F-I) Mean of 3 (G), 4 (B, C, F) or 8 (H, I) independent transfections. Error bars, SEM; *, P<0.05; **, P<0.01; ***, P<0.001; ****, P<0.0001; n.s. not statistically significant; two-way ANOVA with Sidak’s post hoc tests (B, C, F-H) and Student’s t-test (I).

We next examined the effects of the ADD3 KO on Onda-11 proliferation. Consistently with the above, the KO resulted in the opposite phenotypes compared to the OE. The proportions of cells in both G0/early G1 and late G1/S/early G2 phases were reduced, as revealed by both Ki67 expression pattern and EdU treatment (Figure 4D-F and S6A). We further detected an increase in the proportion of cells in G2/M (Figure 4E), but no specific increase in mitotic pVim+ cells (Figure S6B-D).

Finally, we examined if the above effects of ADD3 on cell proliferation are pertinent to U87-MG glioblastoma and H4 neuroglioma cell lines. Similarly to the effects on cell morphology (Figure S3), ADD3 KO only affected the proliferation of U87-MG cells (Figure S7A-C), but not H4 cells (Figure S7D-F). Taken together, ADD3 enables correct Onda-11 proliferation and this effect is relevant also to other glioblastoma cell lines that show morphological heterogeneity.

Considering the dependence of Onda-11 on ADD3 (Figure S1B), we examined the apoptosis of KO cells by immunofluorescence for cleaved caspase-3 (Figure S6B) and detected a marked increase in cell death compared to the control (Figure 4G). Strikingly, this effect was not specific to transfected cells, but we detected a two-fold increase in apoptosis also in the surrounding cells (Figure 4H). Hence, ADD3 is required for the survival of Onda-11 GSCs in both cell-autonomous and non-autonomous manners. Such effects on both the targeted and the neighboring cells prompted us to (1) dissect the changes in the Onda-11 molecular signature following ADD3 manipulation and (2) examine the effects of ADD3 on intercellular connections mediating communication between GSCs.

### Cell-autonomous effects of ADD3 over expression

To elucidate the cell-autonomous effects of ADD3 OE, we performed a bulk RNA sequencing of GFP+ FACS-sorted cell co-transfected with ADD3 or control plasmids. The differential expression analysis revealed 10 upregulated and 7 downregulated genes upon ADD3 OE (Figure 4I). We demonstrated that the genes differentially expressed upon ADD3 OE are indeed exhibiting an expression pattern correlated with ADD3 also at the basal level in other GBM cell lines, i.e., the up-regulated genes are correlated, whereas down-regulated genes are anti-correlated with ADD3 (Figure S8), thus showing robustness of the ADD3 OE signature.

Consistent with the morphoregulatory role of ADD3 (Figure 2), we detected increased expression of cancer-associated palmitoyltransferase *SPTLC3* (Gruel et al., 2014) and secreted protein *SLPI*, involved in filopodia formation (Mizutani et al., 2020). Furthermore, in accordance with the effects of ADD3 OE on GSC proliferation (Figure 4A-C), we detected downregulation of *PLK2*, a key regulator of cell cycle progression, involved in centriole duplication and G1/S transition (Chang et al., 2010; Cizmecioglu et al., 2008).

We next examined whether the effects of ADD3 on cell morphology and proliferation had consequences on cell fate and identity. Since ADD3 induced elongated and branched morphologies of GSCs (Figure 2) and led to a reduction in cell cycle progression and division (Figure 4A-C, S5C) we examined the stemness of ADD3 OE cells and observed that ADD3 sustained as high level of stemness markers as control GSCs (Figure S9A-I). The ADD3 KO in turn led to a minor, albeit not statistically significant, reduction in some of the stemness markers (Figure S9J-R).

Considering that the same morphological and proliferation-related features are also linked to GBM invasiveness (Bhaduri et al., 2020; Venkataramani et al., 2022b), we generated neurospheres from FACS-sorted GFP+ cells over expressing ADD3 or control plasmid and examined their infiltration into the surrounding Matrigel. However, within one week, we did not observe any difference in the invasion index between ADD3 OE and control (Figure S10).

Finally, slowly dividing cells are often associated with therapy resistance (Bao et al., 2006; Chen et al., 2012; Lathia et al., 2015). Interestingly, expression of ADD3 has been previously linked with a population of cells resistant to temozolomide, the main chemotherapeutic used in the GBM treatment (Poon et al., 2015). Furthermore, its expression was also linked to multidrug resistance upon profiling 30 cancer cell lines (Gyorffy et al., 2006). We hence examined the potential signature of chemoresistance among the genes upregulated upon ADD3 OE (Figure 4I) and found *CHI3L1* as a key molecule involved in temozolomide and radio-resistance in GBM cell lines (Akiyama et al., 2014; Shao et al., 2014; Zhao et al., 2020).

Such chemoresistance is strongly associated to a network of tumor-tumor connections (TTCs), including TNTs and TMs (Kolba et al., 2019; Osswald et al., 2015; Wang et al., 2022; Weil et al., 2017). TMs are long protrusions enabling intercellular connections between largely slowly cycling GBM cells (Osswald et al., 2015; Ratliff et al., 2023). Considering that ADD3 promotes protrusion growth and branching along with a reduction in cell cycle progression and that the ADD3 KO has both cell-autonomous and non-autonomous effects on cell survival, we next sought to examine if ADD3-related phenotypes are specifically mediated by TTCs.

### ADD3-induced tumor cell-tumor cell connections (TTCs) are required for the effects of ADD3 on GSC proliferation

To investigate if ADD3 could affect the TTC abundance, we stained Onda-11 GSCs over expressing ADD3 with phalloidin and α-tubulin to detect actin and microtubules, respectively (Figure 5A). We detected doubling of TTCs connecting adjacent cells and containing actin cytoskeleton upon over expression of ADD3 (Figure 5B). Using correlative light-electron microscopy we identified GFP+ co-transfected cells and then examined the ultrastructure of ADD3-induced TTCs using cryo-electron tomography (Figure 5C). This showed that such TTCs are strikingly enriched in actin and that no microtubules were observed. Since the majority of TTCs were short and thin, they were likely TNTs. Nevertheless, we also observed TMs in control Onda-11 (see Figure 1G, H) and upon ADD3 OE (see Figure 5A), which was confirmed by IF for Connexin-43 (Figure S11).

**Figure 5.**
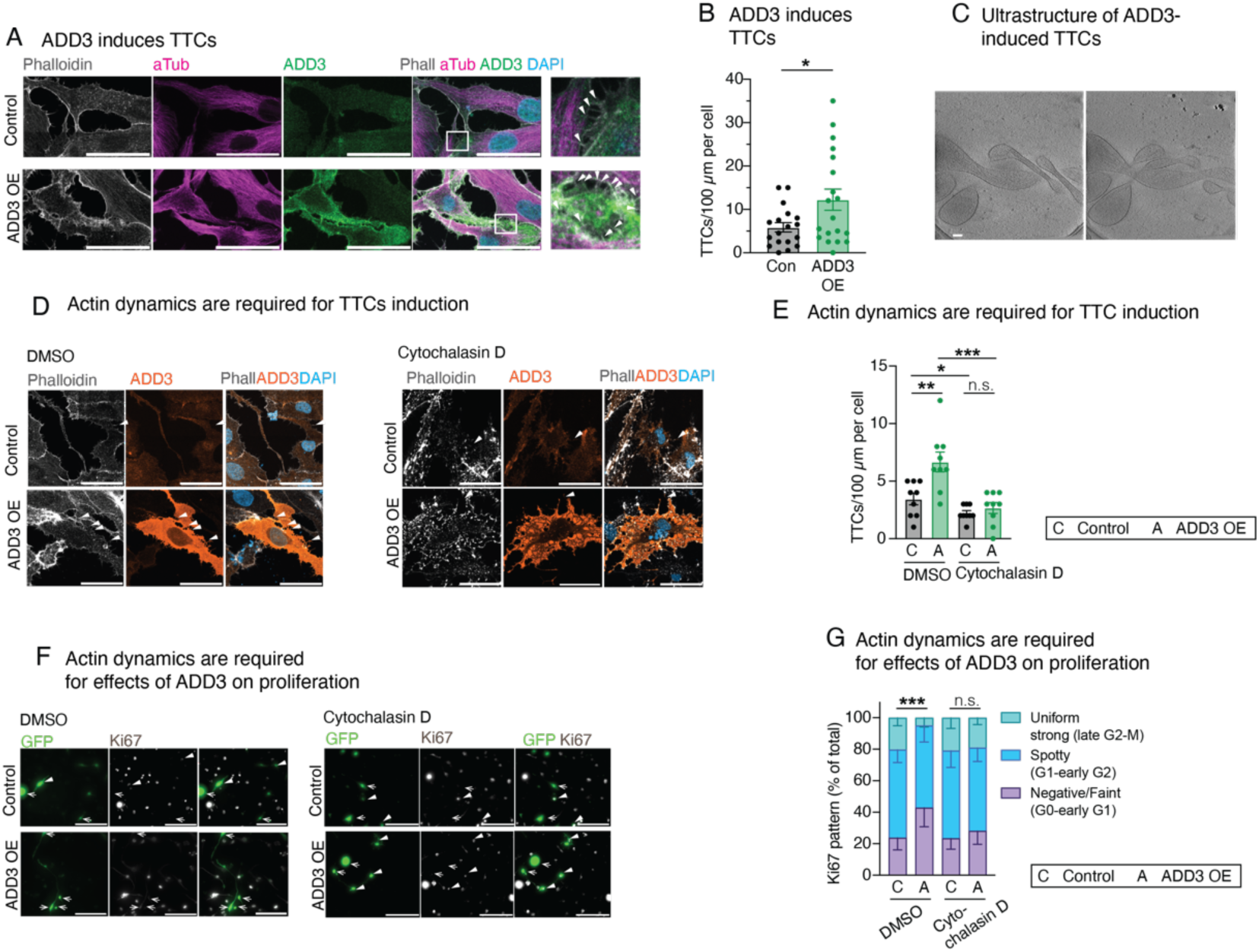
Effects of ADD3 on cell proliferation are mediated by microtubes. (A) IF for actin (Phalloidin, grey), microtubules (α-tubulin, magenta) and ADD3 (green) along with DAPI staining in control (top) and ADD3 overexpressing (OE, bottom) Onda11 GSCs. Arrowheads, microtubes. MIP of 12 planes. Scale bars; 50 µm. (B) Quantification of the number of microtubes per 100 µm of cell perimeter, expressed per cell in control and ADD3 OE. (C) Two slices of a tomogram showing the ultrastructure of ADD3-induced microtubes, extracted from different Z height to show intertwining of the protrusions. Note that the microtubes are rich in actin cytoskeleton. Slice thickness, 10 nm; scale bar, 100 nm. (D-G) Actin cytoskeleton is required for ADD3-mediated both induction of microtubes and effects on proliferation. Following transfection Onda11 GSCs were treated with Cytochalasin D (right) at 5µM concentration for 45 minutes. (D) IF for actin (Phalloidin, grey) and ADD3 (orange) along with DAPI staining in control (top) and ADD3 OE (bottom) Onda11 GSCs treated with 5µM Cytochalasin D (right) and DMSO (left). Arrowheads, microtubes. MIP of 12 planes. Scale bars; 50 µm. (E) Quantification of the number of microtubes per 100 µm of cell perimeter, expressed per cell in control and ADD3 OE upon treatement with 5µM Cytochalasin D or DMSO. (F) IF for Ki67 (white) in GFP+ (green) Onda11 GSCs. Arrows, Ki67 negative/faint; arrowheads, Ki67 spotty/uniform strong. Scale bars, 200µm. (G) Distribution of the three Ki67 patterns of expression in control and ADD3 OE Onda 11 GSCs treated with DMSO and 5µM Cytochalasin. (B, E, G) Mean of 3 independent transfections. Error bars, SEM; *, P<0.05; **, P<0.01; ***, P<0.001; n.s. not statistically significant; Student’s t-test (B, E) and two-way ANOVA with Sidak’s post hoc tests (G).

We thus examined if intact actin cytoskeleton is required for the maintenance of ADD3-induced protrusions by treating the transfected Onda-11 GSC with Cytochalasin D, which causes disruption of actin filaments and inhibits actin polymerization (Figure 5D). Consistently with the above (Figure 5B), DMSO-treated cells exhibited 2-fold increase in TTCs upon ADD3 OE (Figure 5E). In contrast, Cytochalasin D-treated cells lost all the ADD3-induced TTCs and showed similar levels between the control and OE (Figure 5E).

In light of the association of TTCs with cell proliferation (Lu et al., 2019; Osswald et al., 2015; Ratliff et al., 2023; Valdebenito et al., 2018; Venkataramani et al., 2022a) and the ADD3-induced phenotypes on both TTCs and cell cycle progression (Figures 4 and 5A-E), we sought to examine if the effects of the morphoregulatory ADD3 on cell morphology and TTCs are required for its effects on cell proliferation. We treated control and ADD3 OE Onda-11 GSCs with DMSO and Cytochalasin D and examined the expression pattern of Ki67 (Figure 5F), as a key indicator of the effects of ADD3 on cell cycle progression (see Figure 4). Our analysis shows that control cells treated with Cytochalasin D do not have different cell cycle progression compared to DMSO-treated control cells, suggesting that the stability of actin cytoskeleton is not required for their normal proliferation of onda-11 cells. In agreement of what we observed in untreated cells, ADD3 OE GSCs treated with DMSO showed a significant effect on cell proliferation, (Figure 5G and compare to 4B), whereas this effect was completely lost upon treatment with Cytochalasin D (Figure 5G).

Taken together, these data suggest that ADD3 acts as a key regulator of GSC morphology to induce new actin-rich TTCs, which in turn enable cell-cell contacts and mediate the downstream effects on cell proliferation.

## Discussion

In this study we identified the GSC morphology as a key player underlying cell proliferation. We further showed that the main driver of this effect are TTCs. There are three aspects of our study that deserve particular discussion: (1) Cell morphology is a new layer of GBM heterogeneity; (2) GSC morphology affects GSC proliferation and survival through cell-cell connections; (3) ADD3 is a key protein controlling GSC shape.

### Morphology as a new layer of GBM heterogeneity

One of the key reasons for GBM’s malignancy is its extraordinary inter- and intra-tumoral heterogeneity. The molecular heterogeneity, described at genomic, transcriptomic and epigenetic levels, was shown to underlie a multitude of GBM cell types and states (Bhaduri et al., 2020; Chaligne et al., 2021; Couturier et al., 2020; Darmanis et al., 2017; Jacob et al., 2020; Neftel et al., 2019; Patel et al., 2014; Sottoriva et al., 2013). In fact, it has been suggested that each GBM contains on average 11 different cell types (Bhaduri et al., 2020) that could be grouped into four principal cellular states which recapitulate distinct neural cell types (Neftel et al., 2019). Notably, GSCs themselves show a striking molecular heterogeneity within the same tumor (Bhaduri et al., 2020). However, to link these specific cell types with cellular functions and oncological phenotypes, it is also necessary to study potential GBM heterogeneity at the cell biological level.

We have examined GSC morphology and identified four different morphotypes in Onda-11 and U-87 MG cells (Figure 1), suggesting that basic morphological nature is a cell-intrinsic property. The identified morphotypes bore striking similarity to neural stem cells during cortical development, in particular bRG (Kalebic and Huttner, 2020). This is consistent with a large body of evidence showing that GBM initiation, maintenance and progression are controlled by the same signaling pathways and transcription factors that regulate brain development (Azzarelli et al., 2018; Curry and Glasgow, 2021). Despite good molecular understanding, the links between neurodevelopment and GBM at the cell biological level remain largely unexplored. Given that GSCs exist as similar morphotypes to those observed in bRG, where they promote cell proliferation (Kalebic et al., 2019), we asked what the functional consequences of such neurodevelopmental features on GBM are.

A key question to answer was whether the morphoypes are stable or transient cellular states. Our live imaging experiments (Figure 3) suggested that the former is true both within and across cell cycles. It would hence be interesting to examine if such stable morphotypes correspond to transcriptionally defined cell types. So, how is the morphological heterogeneity then generated? Our data showed that the cells of the nonpolar morphoclass are responsible for such heterogeneity, as they are able to generate all the remaining morphoclasses both in mitosis and by morphological transitions in interphase. It is worth noting that nonpolar morphoclass is not present among developmental neural progenitors in interphase (Kalebic and Huttner, 2020). Hence it is tempting to hypothesize that morphological plasticity of nonpolar cells might be a prominent feature of brain cancers, and that it might be linked to the characteristic plasticity among different GBM cell states (Neftel et al., 2019).

### Cell-cell connections link GSC morphology with proliferation and survival

To examine if different morphotypes have distinct cellular functions, we analyzed their proliferation and observed differences in their cell cycle progression (Figure 4). Notably, modifying cell morphology, by over expression of ADD3 and thus generating more elongated cells, led to a reduced cell cycle progression. We found that the key morphological feature responsible for the change in proliferation, is the actin-based tumor-tumor connections (TTCs), including both TNTs and TMs (Figure 5, S10). Whereas formation of TTCs has been implicated in increased cell proliferation (Joseph et al., 2022; Lu et al., 2019; Osswald et al., 2015), a recent study has shown that TM-rich, interconnected GBM cells have a slower cell cycle compared to the fast-dividing, unconnected cells in the invasion zone (Ratliff et al., 2023), which is in agreement with our data (Figures 4 and 5).

Furthermore, such TTC-rich cells over expressing ADD3, did not show altered invasive capacity (Figure S9). This is in line with recent *in vivo* studies showing that a different population of GBM cells, which lacks connections to other GBM cells is the main driver of brain tumor invasion (Ratliff et al., 2023; Venkataramani et al., 2022b). Taken together, our data suggest that TTC-rich GBM cells over expressing ADD3 represent a population of slowly proliferating cells either prior to the infiltration into the brain parenchyma or following it.

TTC-rich GBM cells have also been associated with increased resistance to chemotherapy (Kolba et al., 2019; Osswald et al., 2015; Wang et al., 2022; Weil et al., 2017). Such cells were shown to be able to change their metabolic profile through a TNT-mediated mitochondria, vesicle and protein transfer (Hekmatshoar et al., 2018; Pinto et al., 2021). Interestingly, upon ADD3 OE, we also observed upregulation of CHI3L1, which is involved in chemo- and radio-resistance in GBM (Akiyama et al., 2014; Shao et al., 2014; Zhao et al., 2020), implying that ADD3-induced TTCs might be sustaining therapy resistance. This is further suggested by an increased expression of ADD3 in a population of GBM cells resistant to various chemotherapeutics, including temozolomide (Gyorffy et al., 2006; Poon et al., 2015).

Hence, TTCs appear to be a key feature contributing to the functional consequence of the morphological heterogeneity of GBM. Since GSCs’ transcriptional heterogeneity is mediated by both intrinsic and extrinsic factors (Prasetyanti and Medema, 2017), we propose that the morphological heterogeneity could also be controlled both cell-autonomously and non-autonomously and that TTCs might play a pivotal role in the latter. In fact, we showed that ADD3, as an intrinsic factor promoting morphological heterogeneity, has a critical role in cancer cell survival, both cell-autonomously and non-autonomously (Figure 4G, H). Such effect on surrounding cells might be due to the exchange of specific pro-apoptotic signals from the ADD3 KO cell, or simply because of the lack of any signal from KO cells, following the striking reduction in TTCs.

Taken together, GBM cell morphology mediates intercellular communication and thus has important consequences on tumor cell proliferation, survival, invasiveness and resistance to therapy. Hence, in GBM, like in other cancers (Alizadeh et al., 2020; Barker et al., 2022; Wu et al., 2020), cell morphology has a strong potential to be used as a diagnostic and prognostic marker, through microscopy-based analysis of the tumor.

### ADD3 as a key morphoregulator in GBM

We have identified ADD3 as a key morphoregulator able to control GBM proliferation. We found that ADD3 exerts multiple morphoregulatory functions on GSCs. Notably, it promotes cell elongation and induces various cell protrusions, including TTCs (Figures 2 and 5A-C). Such diverse roles are likely due to its close interaction with actin, a key cytoskeleton component regulating changes in cell shape. Indeed, when the actin cytoskeleton is disrupted, ADD3 is not able to induce TTCs anymore (Figure 5D, E). The question remains if ADD3 directly induces new protrusions by remodeling actin in the membrane cytoskeleton or whether it stabilizes existing protrusions by connecting actin filaments to the plasma membrane. Previous work on other members of the adducin family of proteins seems to favor the latter hypothesis as it has been shown that adducins regulate membrane stability by capping the fast-growing end of actin filaments and connecting spectrin-actin cytoskeleton to membrane proteins (Anong et al., 2009; Baines, 2010; Kuhlman et al., 1996; Li et al., 1998). Accordingly, adducins were shown to stabilize neuronal synapses by controlling spine dynamics (Babic and Zinsmaier, 2011; Bednarek and Caroni, 2011; Pielage et al., 2011). In the context of GBM cells, it is tempting to hypothesize that ADD3 stabilizes cellular projections by providing mechanical support. This ultimately can lead to an increase in the number of stable cell protrusions, particularly long TTCs that enable cell-cell communication.

Considering that such a role of ADD3 on actin cytoskeleton is likely true across different types of cellular projections and cell types, it is plausible that its effects are not specific to Onda-11 GSCs, but generally applicable to GBM cells that are elongated and contain protrusions. In support of this, we showed that ADD3 plays an important role in maintaining the cell morphology of U-87MG cells (Figure S3). Furthermore, ADD3 has already been implicated in GBM progression, therapeutic resistance and cell motility (Kiang et al., 2020; Mariani et al., 2001; Poon et al., 2015; Rani et al., 2013; van den Boom et al., 2003). Nevertheless, the effects of ADD3 on both cell morphology and proliferation are more pronounced in Onda-11 GSC compared to U-87MG (compare Figure 2 to S3 and Figure 4 to S7). We link this to the notion that Onda-11 cells were shown to be strongly dependent on ADD3 in the Cancer DepMap project (Behan et al., 2019; Pacini et al., 2021; Tsherniak et al., 2017), whereas U-87 were not. Beyond the experimental validation of the findings reported in the Cancer DepMap, our results show that DepMap is an important resource for exploring the function of cancer genes in an appropriate model system. In the future it would hence be interesting to study the morphoregulatory mechanisms in more complex model systems such as *in vivo* or in patient-derived organoid system.

Finally, ADD3 was previously shown to regulate the morphology of bRG during brain development (Kalebic et al., 2019). Its KO in human fetal brain tissue led to a reduction in the number of protrusions of neural progenitors, which in turn resulted in a reduction in the proliferative capacity of these cells (Kalebic et al., 2019), which ultimately controls the developmental expansion of the cerebral cortex. This link between cell morphology and proliferation serves as a further example of how neurodevelopment can offer precious insights into brain cancers. It also provides a novel conceptual framework which allows for the identification and mechanistic characterization of other potential molecular targets to be used in future diagnostic and therapeutic approaches in brain cancers.

## Acknowledgements

We are grateful to the services and facilities of HT for the outstanding support provided, notably, N. Maghelli and F. Casagrande from Light Imaging Facility, D. dalle Nogare from Image Analysis Facility, A. Pallini and the team of the Flow cytometry facility, C. Peano and the team of the Genomics Facility and P. Swuec and the team of the CryoEM facility. We thank C. Ossola (Kalebic lab) for the ADD3 clone. We are thankful to B. Soskic (HT) and E. Argenzio (HT) for the critical reading of the manuscript, R. Galli (HSR) for useful comments and all members of the Kalebic lab for helpful discussions. CB and RMI are Ph.D. student within the European School of Molecular Medicine (SEMM). This work has been supported by funds of HT and the grant from AIRC (MFAG 2022 ID 27157) to NK.

## Author contributions

**Carlotta Barelli**: conceptualization, methodology, validation, formal analysis, investigation, data curation, writing-original draft, writing-review and editing, visualization. **Flaminia Kaluthantrige Don**: methodology, formal analysis, investigation, visualization. **Raffaele Iannuzzi**: methodology, formal analysis, visualization. **Ilaria Bertani**: methodology, validation, formal analysis, investigation. **Isabella Osei**: methodology, software. **Simona Sorrentino**: investigation. **Giulia Villa**: investigation. **Viktoria Sokolova**: investigation. **Francesco Iorio**: methodology, writing-review and editing, supervision. **Nereo Kalebic**: conceptualization, formal analysis, writing-original draft, writing-review and editing, visualization, supervision, project administration, funding acquisition.

## Materials and methods

### Reagents and Tools table

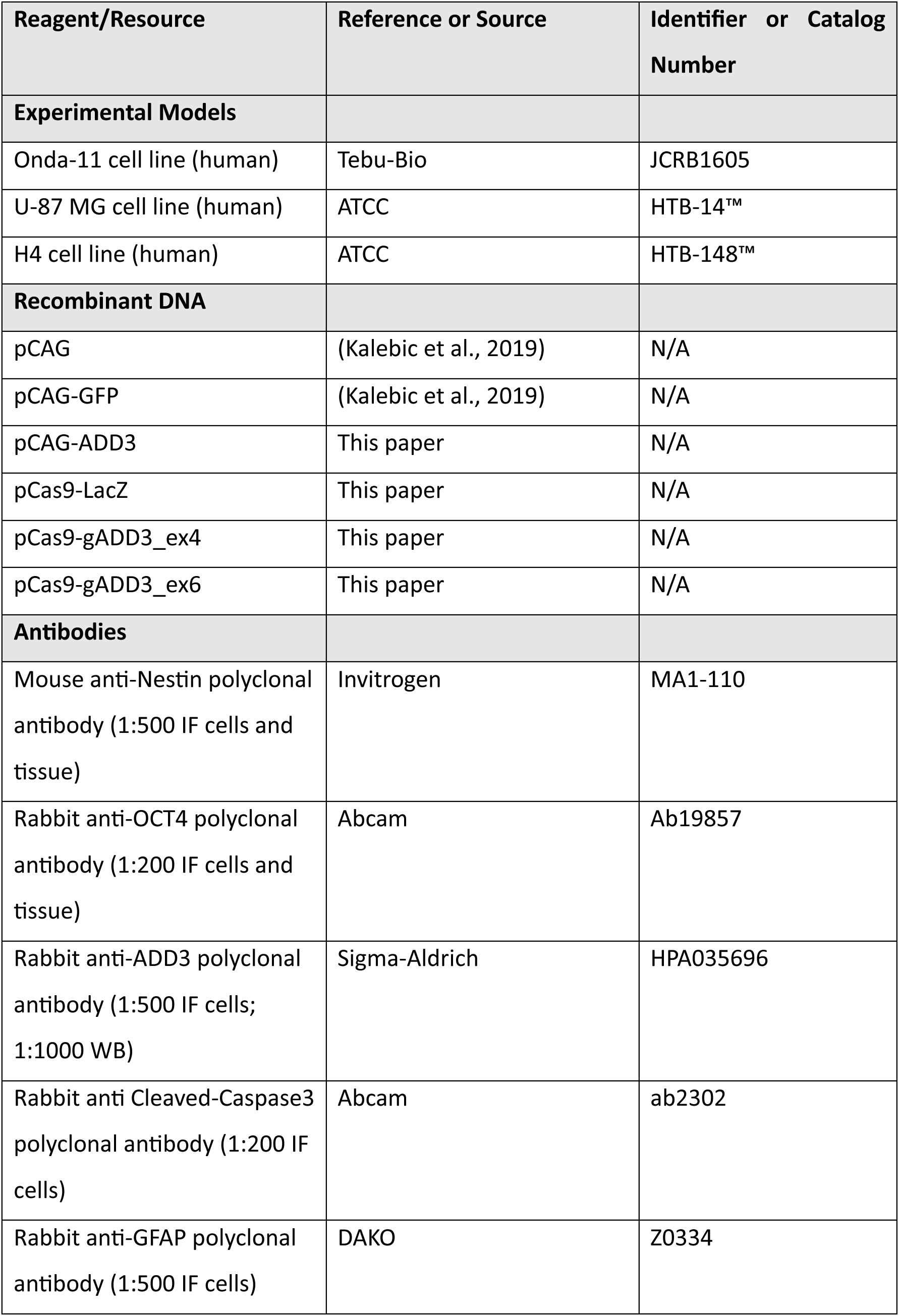
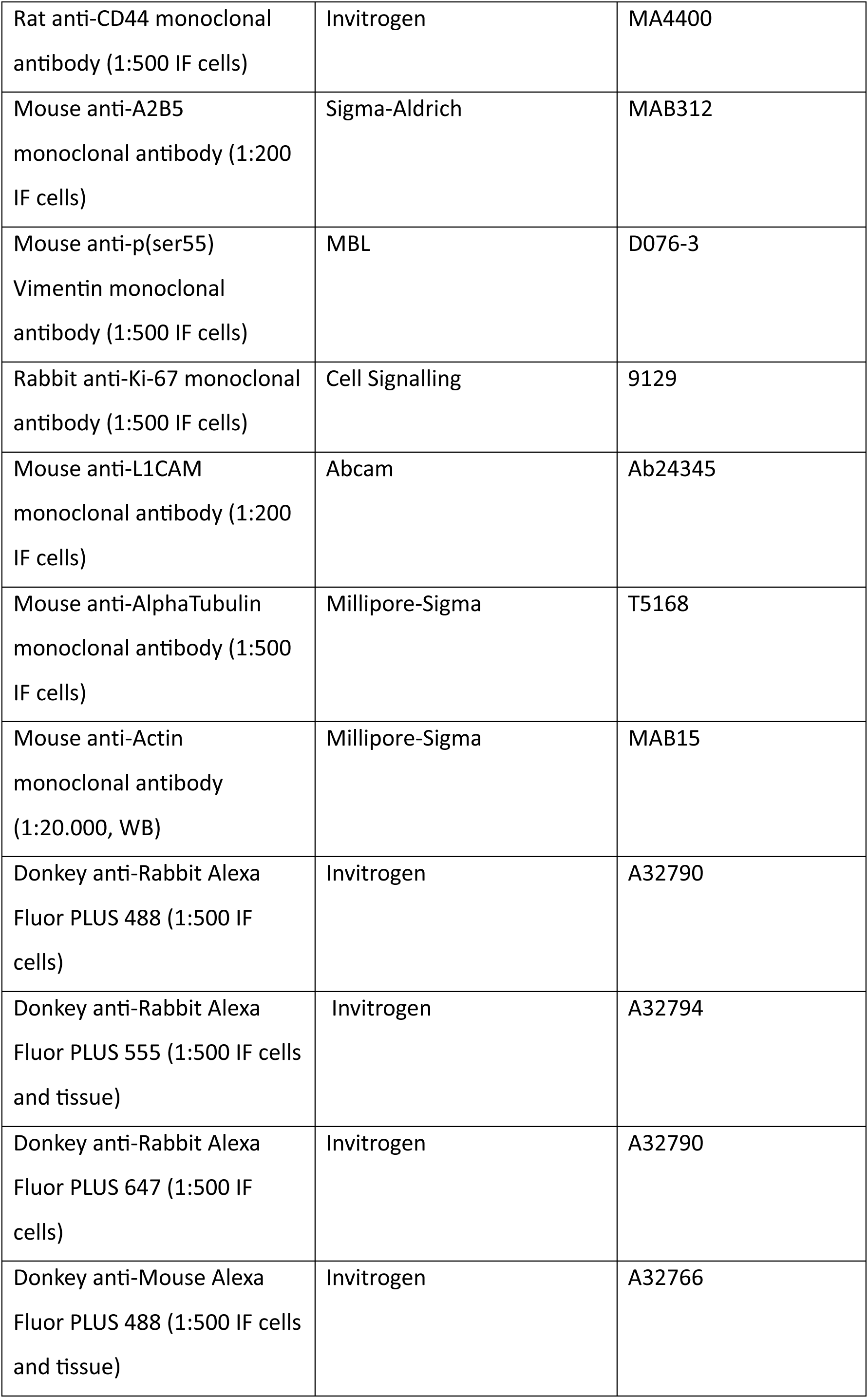
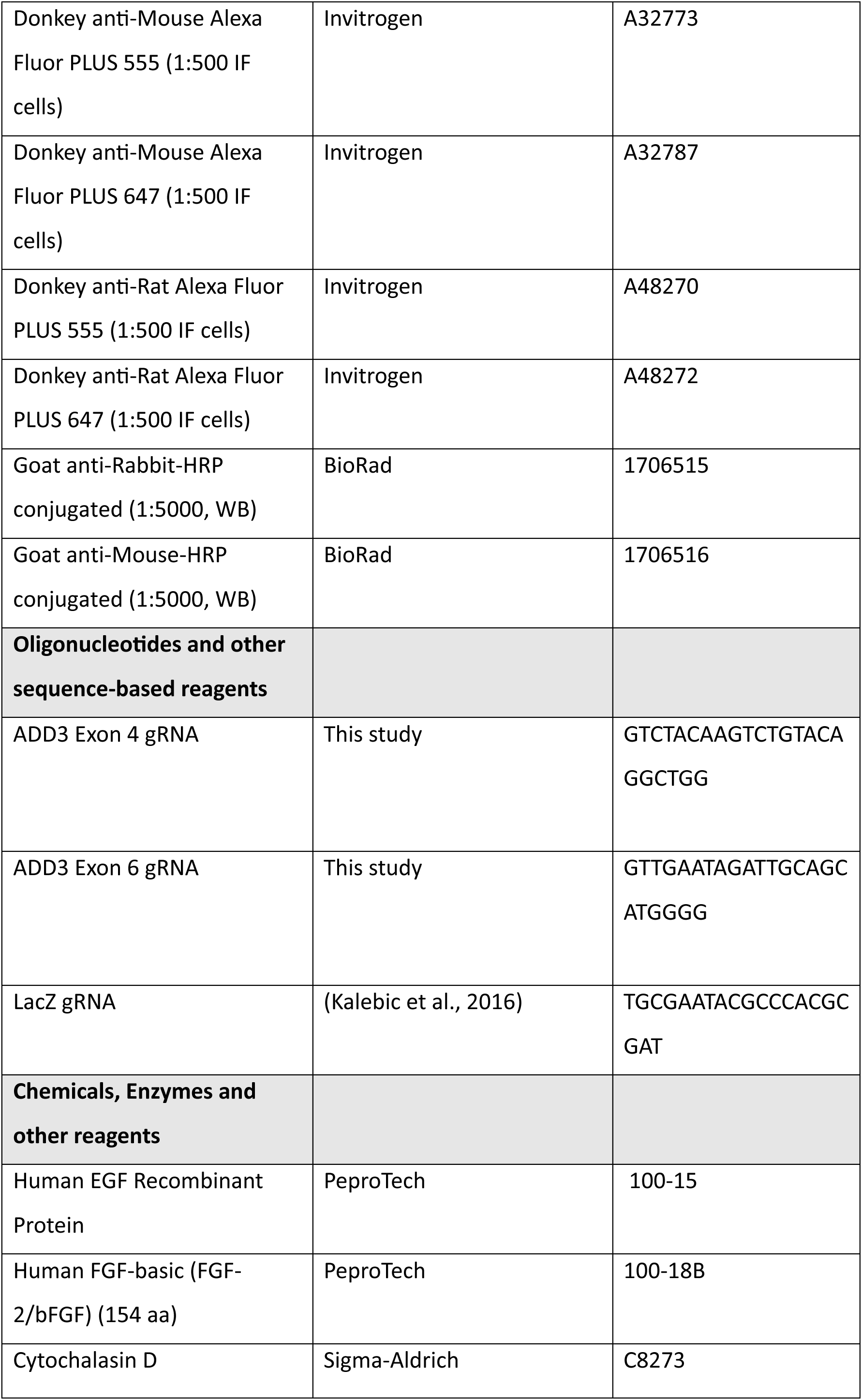
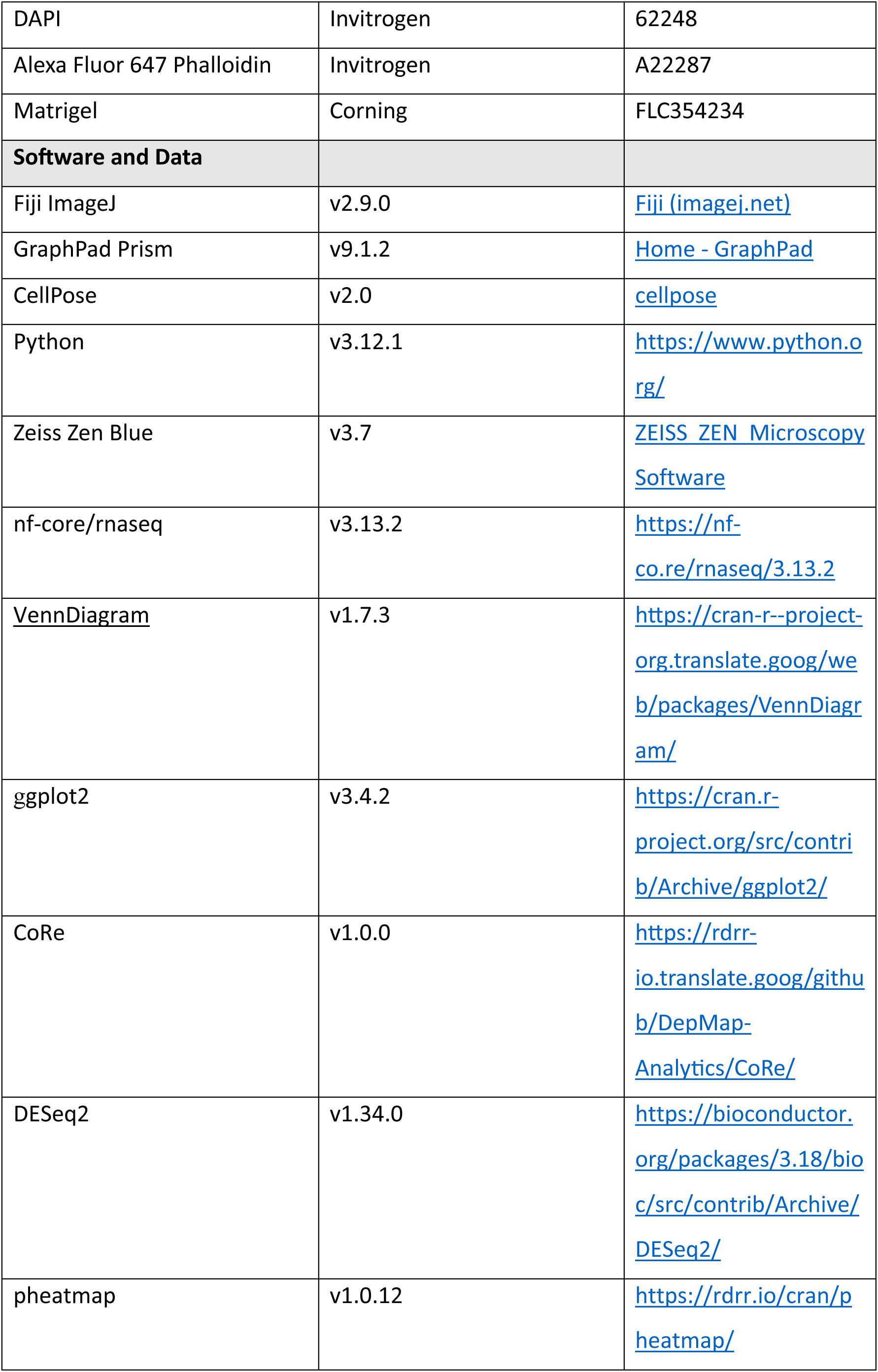
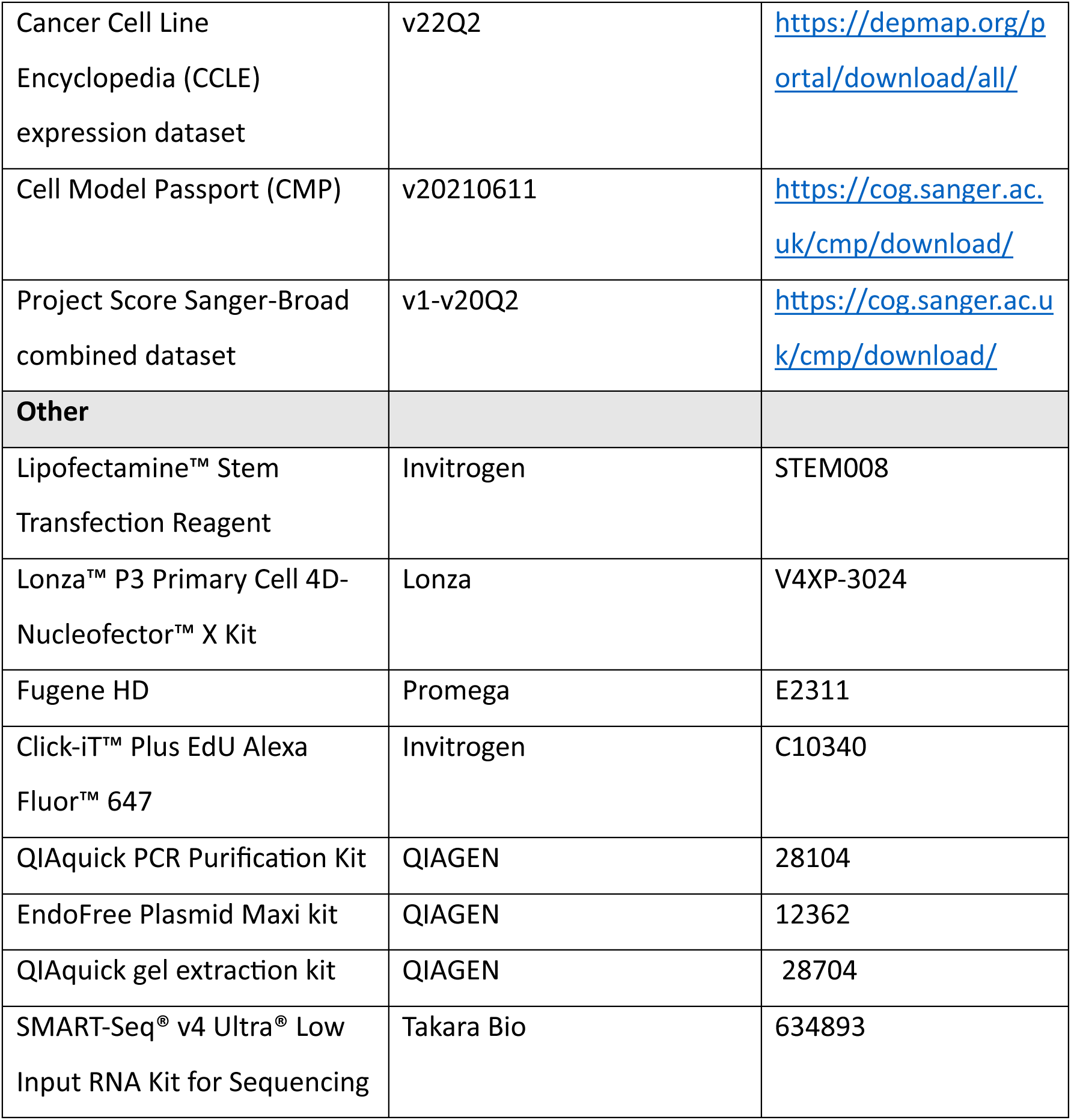

## Methods and Protocols

### Cell culture

Onda 11 cells were reconditioned to glioblastoma stem cells (Onda 11 GSCs) and grown on Laminin(5ug/ml, Sigma, L2020)-coated plates in serum-free media (GSC medium) composed of DMEM/F-12 with 15 mM HEPES and L-Glutamine (Thermo Fisher, 11330057), P/S, N2 supplement (Thermo Fisher, 17502-048), B27 supplement (Thermo Fisher, 17504-044), EGF (10µg/µl) and FGF2 (10µg/µl). U-87MG and H4 cells were grown in DMEM/F-12 with 15 mM HEPES and L-Glutamine, P/S and 10% Fetal Bovine Serum (Sigma, F7524).

For transfection, Onda11 GSCs were plated at a density of 10,000 cells/cm^2^ and treated with OPTIMEM media (Gibco, 31985062) containing Lipofectamine™ Stem Transfection Reagent (LipoStem, Thermo Fisher, STEM008) and DNA mixture. In each six-well plate, 4.5 µl of LipoStem, 2.7 µg of recombinant DNA and 600 µl of OPTIMEM were used. The cells were either fixed after 72 h in 4% PFA for 15 minutes and processed for immunofluorescence (IF), or grown for 48 h, sorted to isolate GFP+ cells and used for RNA and protein extraction, or for neurosphere formation assay. U-87MG cells were transfected with Lonza’s 4D-Nucleofector System following the instructions of the Lonza P3 Primary Cell 4D-Nucleofector X Kit. Briefly, in each cuvette 500,000 cells were treated with 100 µl AMAXA nucleofector solution and 2.5 µg DNA and then re-plated and kept in culture for additional 72-96 h, when they were fixed in 4% PFA. H4 cells were transfected using Fugene HD (Promega, E2311) following supplier’s reverse transfection protocol. Briefly, for each well of a 24 well plate 20 µl of OPTIMEM media, 250 ng DNA and 0.75 µl Fugene HD reagent were washed and 30,000 cells were plated. 72-96h post-transfection the cells were fixed in 4% PFA.

For EdU proliferation assay, 72 h post-transfection Onda 11 GSCs were treated with EdU (Click-iT™ Plus EdU Alexa Fluor™ 647) for 4 h, then fixed in 4% PFA for IF and imaging. For actin cytoskeleton disruption assay, 48 h post-transfection Onda 11 GSCs were treated with Cytochalasin D at a 5 µM concentration for 45 minutes and then kept in culture for additional 4 h and fixed in 4% PFA for IF experiments.

### Cell sorting

The cells were sorted 48 h after transfection to isolate GFP+ cells. MoFLO Astrios EQ cell sorter, equipped with Summit 6.3.1 software (Beckman Coulter), was used for cell sorting prior to the neurosphere formation assay, whereas Cytoflex SRT cell sorter, equipped with CytExpert SRT software (Beckman Coulter) was used for for cell sorting prior to RNA and protein extraction. An average sorting rate of 500–1000 events per second at a sorting pressure of 25 psi (for MoFLO Astrios EQ) or 15 psi (for Cytoflex SRT) with a 100 μm nozzle were maintained.

### Plasmids

For the over expression of ADD3, human *ADD3*–encoding cDNA was amplified by PCR, using the forward and reverse primers CAAX_Xhol_Fw and CAAX_BgIII_Rev as reported above, and cloned into the pCAG vector. DNA was purified using the QIAquick PCR Purification Kit (Qiagen, 28104) and all DNA plasmids were extracted and purified using the EndoFree Plasmid Maxi kit (Qiagen, 12362) following the manufacturer’s instructions.

For CRISPR-Cas9 gene editing of ADD3, two guide RNAs (gRNAs) targeting exons 4 and 6 were cloned into pSpCas9(BB)-2A-GFP (PX458), following the previously published protocol (Ran et al., 2013). For control, a previously published gRNA targeting LacZ was used (Kalebic et al., 2016).

### Immunoblotting

Total cell lysates were prepared in a denaturating buffer (Tris-HCl pH 7.4 50 mM, NaCl 150 mM, 1% SDS). After 15 min of solubilization on a rotating wheel, debris were removed by centrifugation (10,000 g, 15 min at RT). Protein concentration was determined using the Pierce BCA Protein Assay kit (Thermo Fisher Scientific). Total protein extracts (20 µg) were separated on NuPAGE 4–12% gels (Thermo Fisher Scientific) and blotted onto nitrocellulose membranes (Hybond, GE Healthcare). After blocking with 5% dry milk for 1 h at RT, membranes were incubated overnight at 4°C with antibodies against ADD3 (1:1,000) and Actin (1:20,000), washed 3x in TBS-T and incubated with secondary antibodies (1:10,000) for 1 h at RT, washed 3x and the signal was detected using ECL (Clarity western ECL, BioRad) and visualized with a ChemiDoc imaging system (BioRad).

### Neurosphere assay

FACS-sorted GFP+ Onda 11 GSCs were re-plated in ultra-low attachment 96-well plates starting from 1,000 cells per neurosphere in 200 µl GSC medium. Following three days in culture, 50 µl Matrigel was added to each neurosphere. Brightfield images were taken every two days using EVOS M5000 Imaging System with 4X (0.13 NA) or 10X (0.30 NA) objective. Neurospheres were kept till day 15 when they were fixed in 4% PFA for 20 minutes at RT.

### Immunofluorescence (IF)

For IF, the cells were permeabilized for 30 min in blocking solution containing 5% normal donkey serum and 0.3% Triton X-100 in PBS at RT. Primary antibodies were incubated in blocking solution for 2 h, at RT. The following primary antibodies were used: Anti p(ser55)Vimentin (1:500), Anti Ki-67 (1:500), Anti-Nestin (1:500), Anti-ADD3 (1:500), Anti Cleaved-Caspase3 (1:300), Anti-GFAP (1:1000), Anti-CD44 (1:500), Anti-A2B5 (1:300), Anti-L1CAM (1:200), Anti-OCT4 (1:200), Anti-AlphaTubulin (1:500). Following three washes in PBS, the sections were incubated with secondary antibodies (1:500) in 0.3% Triton X-100 in PBS for 30 min at RT, washed again 3 times in PBS and imaged within the following 2 weeks.

### Light microscopy

Confocal microscopy on fixed cells was performed using a Zeiss LSM980 point-scanning confocal or Zeiss LSM980-NLO point-scanning confocal based on Zeiss Observer7 inverted microscopes. The images were acquired with a PlanApo 10X/0.45 dry or a PlanApo 20X/0.8 dry or a PlanApo 40X/1.4NA oil immersion objectives using 405 nm, 488 nm, 561 nm, 639 nm laser lines. The software used for all acquisitions was Zen Blue 3.7 (Zeiss). Once the parameters of acquisition for control conditions had been defined, they were kept constant for all the samples within the same experiment.

Time-lapse imaging on live Onda11 GSCs was performed as follows. 48 h after transfection, the sample was placed under a Zeiss LSM980 point-scanning confocal with a PlanApo 20X/0.8 dry objective and imaged for approximatively 60 h. Z stacks of 18-20 µm range were taken with a Z step of 1 µm and an interval time of 30-40 min.

### Correlative light-electron microscopy and cryo-electron tomography

Quantifoil Gold Grids (R 2/2, Au, 200 mesh, Quantifoil) were plasma-cleaned with a hydrogen and oxygen mix (20:80) for 15 seconds with a Gatan Solarus II and then washed for 1 hour with 100% EtOH. The grids were then coated with 5 ug/ml laminin (for 1 h at 37 °C) and around 25,000 Onda11 GSCs were seeded per grid. 16 h later, the grids were plunged with a Leica EM GP2 plunger. During plunging, a drop of 3 μL BSA-coated 10-nm fiducial gold markers (Aurion) was applied on the EM grids for 2.5 seconds. Grids were stored into liquid nitrogen until acquisition.

Subsequently, cryo-fluorescent imaging was performed on Leica ThunderCryoCLEM system using the Navigator module of Leica LAS X software. Grids were focus-mapped using built-in software functions and imaged in Z-stacks of 10–12 slices and ≈1 µm step size in both transmitted light and green channel fluorescence. The grid maps were saved as .lif files for subsequent identification of the transfected cells at the cryo-transmission electron microscope (cryo-TEM).

Data acquisition was performed using a Thermo Scientific Titan Krios G4 TEM equipped with a Thermo Scientific Selectris X energy filter and a Thermo Scientific Falcon 4i direct electron detector. The microscope was operated at 300 keV in zero-loss mode with an energy filter slit-width set to 10 eV. To identify the area of interest for data collection the map acquired on the Leica Thunder were overlayed with the TEM images acquired with the MAPS (TFS) software. Tomograms were acquired at underfocus from 4 to 6 microns, with a 33K magnification resulting in a 0.376-nm pixel size at the specimen level, using SerialEM software (Mastronarde, 2005). The collection scheme used was dose symmetric, covering an angular range from −60° to +60° with 2° increments, starting at 0°. The cumulative electron dose was ∼120 e−/Å2. All image stacks were motion corrected using alignframes IMOD (Kremer et al., 1996) and reconstructed with AreTomo (Zheng et al., 2022).

### Manual image analysis

All manual cell quantifications were performed in Fiji ImageJ using the CellCounter function, processed with Microsoft Excel, and plotted in GraphPad Prism. For manual analysis of Onda11, U87 and H4 cell morphology, we assigned GFP+ cells to one of the defined morphoclasses and morphotypes. The same was done for Ki67, where GFP+ Ki67+ cells were assigned to one of the three different Ki67 patterns of expression. PVim, Casp3, EdU, L1CAM, A2B5, Nestin, GFAP, OCT4, SOX2 positivity was also calculated using the CellCounter function in Fiji ImageJ. All images were analyzed blindly.

For the time-lapse movies GFP+ morphoclasses were manually tracked overtime and scored for morphological change in interphase and mitosis. Mitotic somal translocation (MST) was defined as the distance the nucleus travels during the time step preceding mitosis. Maximum projections and generations of movies were carried out in Fiji ImageJ.

For neurosphere assay, Image analysis was carried out in Fiji ImageJ where the area of the core and the total neurosphere (including the protrusions) were measured with the freehand line tool. The invasion index was calculated by dividing the area of the core with total area of the neurosphere.

### Automated image analysis

For the machine learning assisted pipeline for image analysis (Figure 2), we collected a total of 39 microscopy images, out of which we segmented the morphology of 1362 Onda11 cells, using CellPose, an artificial neural network for automated cell segmentation. The “cyto2” pretrained model was chosen and retrained for improved Onda 11 cell segmentation. Each cell was labelled through its own image array using Python in a Jupyter notebook. As a first step, each cell was positioned singularly at the center of a new image array with the dimension of the biggest bounding box and saved as ‘tiff’ file. Subsequently, the following morphological features were extracted: area, perimeter, major and minor axis lengths, eccentricity. These properties were engineered using the “regionprops_table” function from the scikit-image library to compute properties (measurements) out of labelled regions in the image arrays. Eccentricity is a measure of cellular elongation and circularity, where an eccentricity equal to 0 indicates a circle, whereas values between 0 and 1 indicate an ellipse.

To analyse Onda 11 cell protrusions, we modified our previous semi-manual workflow named Progenitors Process Analysis (PPA) (Kalebic et al., 2019) and used to quantify number of primary and all protrusions, average and maximum protrusion length, branching index (ratio between the total number of protrusion and primary protrusions) and Sholl analysis.

### Data driven selection of ADD3

To identify genes that potentially regulate glioblastoma stem cell morphology, we used a published tumor atlas of differentially expressed genes in primary glioblastoma tumors (Bhaduri et al., 2020). We intersected this dataset with a list of morpho-regulatory genes involved in neurodevelopment identified in (Kalebic et al., 2019). This yielded a list of 30 candidate genes. The enrichment of adducins among the 30 genes was calculated using the following parameters: total number of human protein coding genes = 19,396 (N), total number of adducins = 3 (n), number of selected genes = 30 (k), number of hits = 3 (x). We then investigated the expression level of the selected genes (29/30 genes as one of the genes, MGEA5, was not analyzed in the datasets mentioned below) in 48 annotated glioblastoma cell lines from Cancer Dependency Map Dataset (22Q2 version) (Behan et al., 2019; Pacini et al., 2021; Tsherniak et al., 2017) and the Sanger Cell Model Passport (van der Meer et al., 2019) observing a bimodal distribution from which we identified 18 highly expressed genes (whose basal expression was seemingly generated by the distribution with the higher mean). Subsequently, we derived the depletion fold change of these 18 genes upon CRISPR-Cas9 targeting in 48 GBM cell lines using the same resources. We excluded pan-cancer core fitness genes (as predicted in (Vinceti et al., 2021)) and focused our attention on ADD3 as an important morphoregulator during development (Kalebic et al., 2019), differentially expressed in GBM (Bhaduri et al., 2020) and with a strong and context specific depletion fold change in GBM cell lines. We then identified Onda 11 as the GBM cell line with the highest dependency on ADD3. U-87 MG was selected as a GBM cell line with low or no ADD3 dependency, while H4 were selected as glioma cell line with mild ADD3 dependency.

### RNA sequencing and gene expression analysis

During sorting, GFP+ Onda 11 GSCs were collected in lysis buffer containing RNA inhibitor in nuclease-free water. RNA was extracted through the SMART-Seq® v4 Ultra® Low Input RNA Kit for Sequencing (Takara). The libraries were sequenced with NovaSeq 6000 with SP flow cell and the following read configuration: 150×10×10×150. Reads from the same sample, obtained from different sequencing lanes, were aggregated and subjected to adapter trimming using Trim Galore. Processed reads were aligned to the human reference genome (GRCh38) using STAR and quantification performed with Salmon. Count data were regularized and log transformed using the *rld* built-in DESeq2 function and samples were clustered based on Euclidean distances. Differential expression analysis was performed using DESeq2 using raw counts as input. Differentially expressed genes were identified using a cutoff of absolute log_2_ fold change (log_2_ FC) ≥ 0.5 and False Discovery Rate (FDR) < 0.05. To comprehensively evaluate the outcomes of the Differential Expression Analysis, we employed Cancer Cell Line Encyclopedia (CCLE) profiles - standardized to achieve zero mean and unit variance – across 48 GBM cell lines. We calculated pair-wise correlation scores across all genes, considering the upper triangle of this matrix as a null distribution of scores. Pair-wise Pearson’s correlation scores between ADD3 and DEGs were extracted and compared to the null with a t-test.

### Statistical analysis

All statistics analyses were conducted using Prism (GraphPad Software). To test for statistical significance (p<0.05), two-way ANOVA with Sidak or Bonferroni post hoc tests, Fisher exact test and Student’s t test were used. For each graph, the number of samples, statistical test and the p value are noted in the figure legends.

